# Copy number variant syndromes are frequent in schizophrenia: progressing towards a CNV-schizophrenia model

**DOI:** 10.1101/596213

**Authors:** Venuja Sriretnakumar, Clement C. Zai, Syed Wasim, Brianna Barsanti-Innes, James L. Kennedy, Joyce So

## Abstract

The genetic underpinnings of schizophrenia (SCZ) remain unclear. SCZ genetic studies thus far have only identified numerous single nucleotide polymorphisms with small effect sizes and a handful of copy number variants (CNVs). This study investigates the prevalence of well-characterized CNV syndromes and candidate CNVs within a cohort of 348 SCZ patients, and explores correlations to their phenotypic findings. There was an enrichment of syndromic CNVs in the cohort, as well as brain-related and immune pathway genes within the detected CNVs. SCZ patients with brain-related CNVs had increased CNV burden, neurodevelopmental features, and types of hallucinations. Based on these results, we propose a CNV-SCZ model wherein specific phenotypic profiles should be prioritized for CNV screening within the SCZ patient population.

## 1. INTRODUCTION

Schizophrenia (SCZ) has a high heritability of up to 0.85 but, despite numerous large-scale genetic studies, only genetic variants with primarily small effect sizes have been identified to date (Allen et al., 2008; Bergen and Petryshen, 2012; Craddock et al., 2005; Owen et al., 2003; Ripke et al., 2014; Sebat et al., 2009; Shih et al., 2004). Chromosomal copy number variants (CNVs) have been found to contribute more significantly to the risk of SCZ than most other genetic variants (Zhuo et al., 2017). To date, several recurrent microdeletions and microduplications have been implicated in SCZ, the most well-known being microdeletion 22q11.2 syndrome, also known as DiGeorge or velocardiofacial syndrome (DGS)(Bassett et al., 2008; Karayiorgou et al., 2010). Recent epidemiological studies show that up to 1% of all SCZ patients harbour the DGS deletion, while 22-30% of all individuals with DGS develop SCZ or schizoaffective disorder, a prevalence of about 25 to 30 times higher than the general population (Bassett and Chow, 2008). Though there have been many large-scale studies investigating recurrent CNVs unique to the SCZ patient population, with the exception of DGS, there is a lack of scientific literature exploring the prevalence of other already known and clinically well-characterized syndromic CNVs that could be playing a role within the SCZ population (Consortium, 2008; Marshall et al., 2017; Stefansson et al., 2008). Examples of syndromic CNVs that have been associated with SCZ include microdeletions at 1q21.1, 15q13.3, 16p12.2 and 22q13.3, and microduplications at 1q21.1 and 17q12 (Consortium, 2008; Forsingdal et al., 2016; Nevado et al., 2014; Phelan and McDermid, 2011; Rasmussen et al., 2016). Interestingly, most of these syndromic CNVs related to SCZ are also concurrently associated with other brain-related phenotypes, including autism spectrum disorder (ASD), bipolar disorder and intellectual disability (ID)(Chen et al., 2016; Deshpande and Weiss, 2018; Forsingdal et al., 2016; Larson et al., 2018). This is in line with the neurodevelopmental continuum model, wherein SCZ forms part of a spectrum of neurodevelopmental disorders, including ASD, bipolar disorder, ID and attention-deficit/hyperactivity disorder, suggesting common underlying pathogenetic pathways (Davis et al., 2016; Owen and O’Donovan, 2017).

In this study, we explore the contribution of known syndromic CNVs and candidate CNVs that may underlie the phenotypes seen in a large SCZ cohort.

## 2. MATERIALS AND METHODS

### 2.1 Samples

DNA samples from 348 SCZ patients at the Centre for Addiction and Mental Health (CAMH, Toronto, Canada) were genetically analyzed in this study. The sample characteristics, a subset of which is investigated in this study, were previously described in (Zai et al., 2010). Briefly, the sample is comprised of primarily Caucasian ancestry, with an average age of 49.8 at the time of recruitment. The detailed demographic characteristics of the sample is shown in Supplementary Table S1. Consent was obtained at the time of recruitment and the study was approved by the CAMH research ethics board.

### 2.2 CNV Detection

CNVs were detected using the Affymetrix Genome-Wide Human SNP Array 6.0 (Cat No. 901182, ThermoFisher Scientific Inc., Waltham, MA, USA) following the manufacturer’s protocol.

### 2.3 Phenotyping

A retrospective review of the Structured Clinical Interview for DSM-IV (SCID) and medical records available at time of study recruitment was conducted to collect data regarding height and weight, psychiatric and systemic medical co-morbidities, neurodevelopmental and neurological findings, types of hallucinations, treatment response, and family history. Phenotypic information was coded using the International Classification of Disease 10 (ICD-10 Version: 2016) for further analysis.

### 2.4 CNV Calling

CNVs were called using Genotyping Console™ (GTC) and PennCNV software (Wang et al., 2007) using HapMap reference samples. In order to increase accuracy and decrease false positive CNV calls, BEDTools v.2.27.0 was used to filter for overlapping CNVs called by both software, which were then used for all further analyses (Quinlan and Hall, 2010; Zhang et al., 2014). Size thresholds of 200 kb and 500 kb were applied for deletions and duplications, respectively.

### 2.5 Cluster Analysis

Cluster analysis was performed using R language version 3.4.3 to empirically determine whether the patient CNV and phenotype dataset contained any distinct groups/clusters (Supplementary Figure S1)(Team, 2017).

### 2.6 CNV Gene Analysis

CNVs were cross-referenced against the Human Genome build GRCh37/hg19 for overlapping OMIM and RefSeq genes using the UCSC Genome Browser(Kent et al., 2002; McKusick-Nathans Institute of Genetic Medicine, 2018; O’Leary et al., 2016). Gene enrichment analysis was conducted on all genes within the CNVs using Gene Ontology (GO) Panther 13.1 gene list analysis (Thomas et al., 2003). Gene enrichment was replicated using the Reactome Pathway Browser 3.4 (Croft et al., 2014; Fabregat et al., 2018). FUMA GENE2FUNC was used to determine the tissue specificity of the genes (Watanabe et al., 2017). All tests were run using default settings.

### 2.7 Characterization of Syndromic and Candidate Brain CNVs

CNVs were cross-referenced to clinically well-characterized syndromic CNVs that are known to present with neuropsychiatric features, such as a number of those listed in NCBI Gene Reviews^®^ and by search of the medical literature using PubMed. Patients identified to carry a syndromic CNV were included in the “known brain CNV” group for further analysis. All identified genes within the CNVs were manually curated for association with neurological, neurodevelopmental and/or neuropsychiatric phenotypes. Patients with CNVs containing genes associated with these brain-related phenotypes who were not already part of the “known brain CNV” group were categorized into the “candidate brain CNV” group. All remaining patients without “known brain” or “candidate brain” CNVs were included in the “no brain CNV”group.

### 2.8 Statistical Analysis

Various characteristics amongst the CNV groups were statistically compared using Mann Whitney U test for all scalar data and Chi-squared test for all categorical data. For all significant Chi-squared test results, *post hoc* group-by-group analyses were conducted to determine which groups were statistically significant. All significant p-values were corrected using False Discovery Rate (FDR) for the total number of tests done. Binary logistic regression was carried out to determine if the significant variables were reliably able to predict the types of CNV groups. All tests were performed using R language version 3.4.3 and/or IBM SPSS Statistics for Windows Version 20.0 (Team, 2017).

## 3. RESULTS

### 3.1 CNV Calling

A total of 1 032 CNVs, including 861 deletions and 171 duplications, were identified (Supplementary Figure S2).

### 3.2 Cluster Analysis

K-modes cluster analysis did not yield any insights into the structure or classifications of phenotypes relative to CNVs (Supplementary Figure S1).

### 3.3 CNV Gene Analysis

GO Panther and Reactome Pathway analyses resulted in consensus findings showing an enrichment of genes within immune system pathways (Supplementary Table S2). Specifically, gene enrichment was seen for the adaptive immune system and cytokine signaling in immune system pathways. FUMA GENE2FUNC illustrated several differentially expressed gene sets for various tissue types, including various brain regions, for genes from the CNVs called compared to a list of background genes (Figure1).

**Figure 1.**
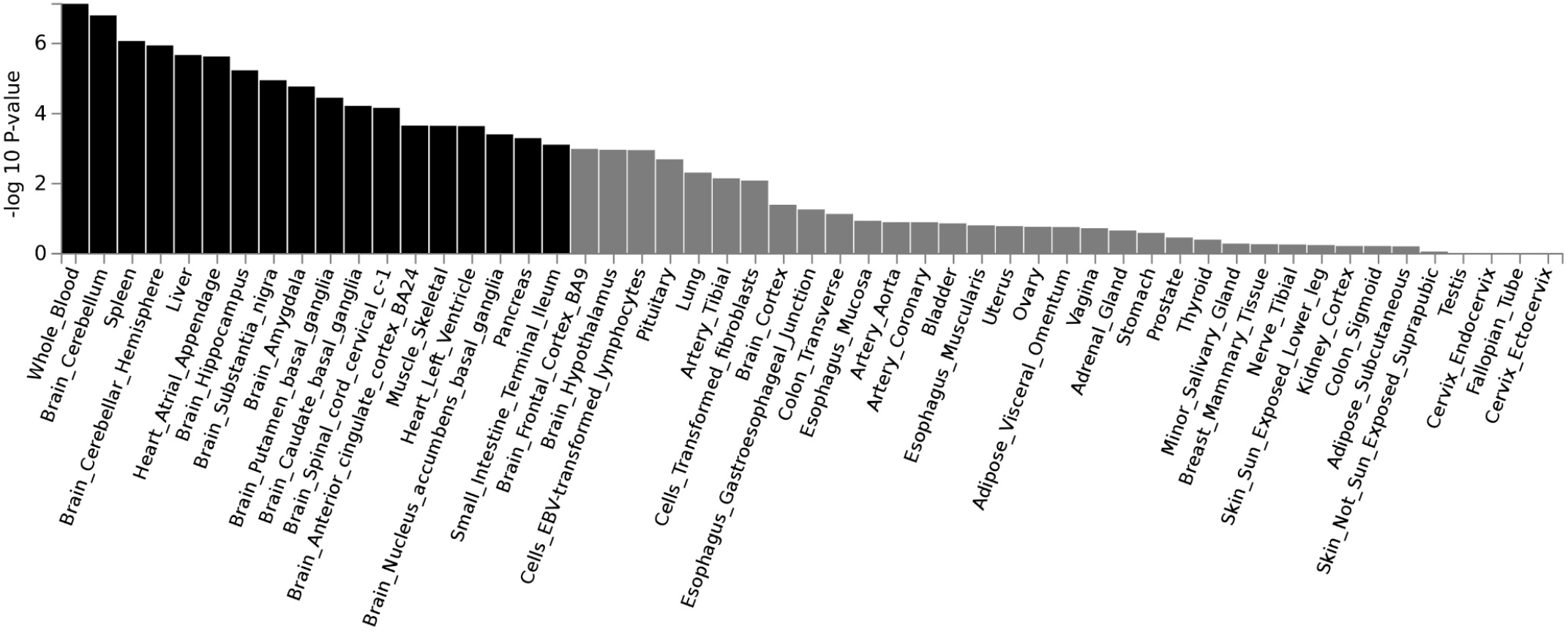
Tissue specificity graph of differentially expressed gene (DEG) set from all CNVs called. Significantly enriched DEG sets (P_bon_ < 0.05) are depicted by the black bars. Image adapted from FUMA GENE2FUNC.

### 3.4 CNV Characterization

A breakdown of the number and types of CNVs is shown in Supplementary Figure S2. Fourteen patients were identified with a total of 12 distinct CNVs associated with previously recognized syndromic CNVs (“known brain CNV” group; Table 1, Supplementary Table S3). A further 30 patients were identified to carry CNVs containing or disrupting one or more genes associated with brain-related phenotypes (“candidate brain CNV” group; Table 1, Supplementary Table S3 and S4). Eight patients in the “known brain CNV” group also carried additional CNVs containing candidate genes of interest. The remaining 304 patients were categorized into the “no brain CNV” group. Taken together, in our SCZ cohort, 4.02% (14/348) of patients were identified to have known pathogenic CNVs and 8.62% (30/348) have candidate brain-related CNVs, for a combined total of 12.64% (44/348) of SCZ patients with brain-related CNVs of interest.

**Table 1.** Patient CNV and phenotype data.

| ID | Cytoband | Known/<br>Candidate<br>SCZ genes | Phenotypic Findings |  |  |  |  |  |  |  |  |
| --- | --- | --- | --- | --- | --- | --- | --- | --- | --- | --- | --- |
|  |  |  | DSM-5 |  | Hall | TRx | Neuro | Systemic | Psychiatric<br>Family Hx |  | Other FHx |
|  |  |  | DD/ID | Other |  |  |  |  | Psych | Other |  |
| 15 | Del 1q21.1 | *PEX11B* |  | DEP, PD | A, V | + |  |  | + | + | ID, PD |
| 1* | Del 1q21.1-q21.2 | *GJA8* |  |  |  |  |  |  |  |  |  |
|  | **Del 16p11.2** | -- |  |  | A, G/O, V | - |  |  | - | + | Ep |
|  | Del 22q11.21 | *COMT* |  |  |  |  |  |  |  |  |  |
| 21 | Del 2p16.3 | *NRXN1* | + | PD | A, V | - |  | O | - | + |  |
| 22* | Del 2p16.3 | *NRXN1* | + | CD | A, V | + |  | O, HN | - | + | ID; DM |
| 10* | **Del 5q35.3** | *NSD1* |  |  | A, G/O, V | - |  | DFS | - | + | Glossodyn |
| 11 | Del 15q11.1-q12 | -- |  | DEP | A | - |  | T2DM | - | - |  |
| 12* | **Del 16p11.2** | *ALDOA, TBX6* |  |  | A, G/O, T | + | Ep, Ab-Brain-EEG, Narcolepsy |  | - | + |  |
| 13* | **Del 16p11.2** | *ALDOA, TBX6* |  |  |  |  |  |  |  |  |  |
|  | **Del 16p13.3** | *TSC2* |  | DEP, SWD | A, T, V | + |  | O, T2DM | - | + | SIDS |
| 4 | Del 16p13.11-p12.3 | *ABCC6* |  |  | A, G/O, V | - |  |  | - | - |  |
| 14 | Dup 16p13.11-p12.3 | *ABCC6* |  | DEP | A | - |  | DM | - | - |  |
| 2* | **Del 17p12** | *PMP22* |  |  | A, G/O, T, V | + |  | O | + | + | LD |
|  | Del 22q11.21 | *COMT* |  |  |  |  |  |  |  |  |  |
| 3 | **Del 17p12** | *PMP22* | + | ADHD | A, T, V | + | PN |  | - | + | PN |
| 37* | **Dup Xp22.12** | *CNKSR2* | + | ANX, CD, DEP, PD, SD, TD, | A, T, V | + |  | O | + | + | Ep; SIDS |
| 44*† | **Dup Xp22.12** | *CNKSR2* |  |  |  |  |  |  |  |  |  |
| 23 | Del 2q11.2 | *LMAN2L* |  |  | A, G/O, V | - |  | O | - | - |  |
| 40 | Del 2q11.2 | *LMAN2L* |  | ANX | A, V | + |  |  | - | - | DM |
| 42 | Del 2q11.2 | *LMAN2L* |  |  | A, T | + |  | O | - | + |  |
| 32* | Del 6p21.33 | *HLA-B, MICB* |  |  |  |  |  |  |  |  |  |
| 39 | Del 5p12 | *HCN1* |  | DEP, SS | A, T, V | + |  |  | + | + |  |
| 33* | Del 5p12 | *HCN1* |  |  | V | + |  | O | + | - |  |
| 41* | Del 5p12 | *HCN1* |  |  | A | + |  |  | + | - |  |
| 30* | Del 5p12 | *HCN1* |  |  | A | + |  |  | - | - |  |
| 29* | Del 5p12 | *HCN1* |  | SS | A | + |  | Psoriasis | - | + |  |
| 5 | Del 16p13.11 | *ABCC6* | + | DEP | A, T, V | + | Ab-Brain-Imaging |  | - | + |  |
| 6 | Del 16p13.11 | *ABCC6* |  |  | A | - |  |  | + | + |  |
| 7 | Del 22q11.21 | *COMT* |  |  | A, V | + |  |  | + | + |  |
| 8 | Del 22q11.21 | *COMT* |  |  | A, G/O, V | + |  |  | - | + |  |
| 9 | Del 22q11.21 | *COMT* |  | DEP | - | + |  | DM | - | + |  |

### 3.5 Statistical Analysis

Phenotype comparisons of the brain-related CNV groups (Table 2) revealed that the “known brain CNV” group had significantly increased numbers of total CNVs and neurodevelopmental phenotypes (i.e. developmental delays, ID), and was significantly more likely to present with more than two types of hallucinations (i.e. auditory, visual, tactile, olfactory/gustatory) compared to the “no brain CNV” group. The “candidate brain CNV” group had a significantly higher number of total CNVs and greater prevalence of tardive dyskinesia (TD) compared to the “no brain CNV”group. No significant differences were observed between the “known brain” and “candidate brain” CNV groups (data not shown), with the exception of neurodevelopmental phenotypes.

**Table 2.** Statistical comparison of the “known brain”, “candidate brain”, and “no brain” CNV groups of SCZ patients.

|  | "known brain" | "candidate brain" | "no brain CNV" |
| --- | --- | --- | --- |
| Avg. # of CNVs per patient (±S.E.M.) | **7.08±1.47** | **6.57±1.06** | 2.47±0.15 |
| Neurodevelopment phenotype (%) † | **5/10 (50.00)** | 1/25 (4.00) | 10/216 (4.63) |
| Tardive dyskinesia phenotype (%) | 0/10 (0.00) | **6/25 (24.00)** | 18/216 (8.33) |
| Avg. number of types of hallucinations (±S.E.M.) | **2.46±0.29** | 1.97±0.18 | 1.73±0.07 |

*Statistically significant differences ( $p$ -value<0.05) compared to the “no brain CNV” group are bolded. † There is a significant difference between the prevalence of neurodevelopmental phenotypes between “known” and “candidate brain” groups, $P_{FDR}$ <0.01. Avg, average; CNV, copy number variants; S.E.M., standard error of mean.*

Because enrichment of immune system-related genes was identified in the CNVs of the study cohort, similar phenotype comparisons were performed between patients with (126/348) and without (222/348) immune-related CNVs. Patients carrying CNVs with immune pathway genes were categorized in the “immune CNV” group, while those without CNVs containing immune-related genes were placed in the “no immune CNV” group. The total number of CNVs per patient was significantly higher in the “immune CNV” group, as were the number of patients with brain-related CNVs, average number of brain-related CNVs per patient, and the proportion of brain-related CNVs to total CNVs per patient in comparison to the “no immune CNV” group (Table 3). No significant phenotypic correlations were seen.

**Table 3.** Statistical comparison of “immune” and “no immune” CNV groups of SCZ patients.

|  | "immune" | "no immune" |
| --- | --- | --- |
| Avg. # of CNVs per patient (±S.E.M.) | **8.00±1.60** | 1.92±0.13 |
| # of patients with brain CNVs (%) | **26/126 (20.63)** | 18/222 (8.11) |
| Avg. # of brain CNVs per patient (±S.E.M.) | **0.39±0.09** | 0.10±0.02 |
| Proportion of brain CNVs to total CNVs per patient (±S.E.M.) | **0.05±0.01** | 0.03±0.01 |

*Statistically significant differences ( $p$ -value $<0.05$ ) compared to the “no immune” CNV group are bolded. Avg, average; CNV, copy number variants; S.E.M., standard error of mean.*

Binary logistic regression analysis found that presence of neurodevelopmental phenotypes and number of immune genes per patient were significant predictive factors for the “known brain CNV” group (χ^2^=30.01, df=2, p<0.001). This model explained 39.6% (Nagelkerke R^2^) of variance and correctly classified 97.2% of patients with known syndromic CNVs. Specifically, increasing number of immune genes within CNVs (OR=1.70, 95% C.I. 1.09 – 2.65) and presence of neurodevelopmental phenotypes (OR=18.88, 95% C.I. 3.89 – 90.91) were associated with an increased likelihood of harboring known syndromic CNVs. Additionally, individuals with more than two types of hallucinations were at five times the odds (95% C.I. 1.57 – 15.63) of having known syndromic CNVs and those with TD were 3.6 times more likely to have candidate brain CNVs.

## 4. DISCUSSION

In our study of CNVs in 348 SCZ patients, we found a significant proportion of patients with known (4.02%) and candidate (8.62%) brain-related CNVs. Studies thus far have been unable to establish the exact contribution of CNVs to the risk of development of SCZ due to sample size limitations. Walsh et al. (2008) found that 15% of their cohort of 150 SCZ patients carried CNVs disrupting genes in neurodevelopmental pathways. Meanwhile, it is estimated that CNVs contribute to approximately 10% of neurodevelopmental disorder cases (Williams et al., 2009), with some studies reporting even higher rates (e.g. 13% pathogenic CNVs in ID patients with comorbid psychiatric features, 14.3% pathogenic CNVs in complex ASD patients)(Lovrecic et al., 2018; Thygesen et al., 2018). This is comparable to the combined prevalence of patients in our study with known and candidate brain-related CNVs (12.64%). Moreover, ten patients from our study carry CNVs affecting eight genes (*ALDOA, CNKSR2, CNTN4, DOC2A, HCN1, HIRIP3, SEZ6L2*, and *TBX6*) that overlap with genetic risk loci reported from the largest SCZ genome-wide association study to date (Ripke et al., 2014). Interestingly, amongst patients found to have brain-related CNVs, nearly a third (14/44) have one or more additional brain-related CNV, and there was a significantly increased number of total CNVs (i.e. CNV burden) compared to those without brain-related CNVs. This is in line with the concept that the high heritability of SCZ is conferred by a large number of genetic risk factors with small effect sizes (Ripke et al., 2014). A recent large-scale study (Marshall et al., 2017) showed increased CNV burden amongst 21 094 SCZ patients compared to 20 227 controls, and the CNVs in the SCZ cohort were significantly enriched for genes associated with neurobehavioral phenotypes and synaptic functions. Together, these results emphasize the importance of further studies to determine whether CNV burden modulates phenotypes in SCZ patients.

Our study was limited by sample size. Due to the rarity of individual CNVs, research of this nature requires large sample sizes to be sufficiently powered to detect significant differences. Post-hoc power analysis was conducted for each family of tests and comparisons and is summarized in Supplementary Table 5. Overall, sufficient power was achieved for primary comparisons against control. Based on the results of this study, appropriate sample size calculations can be conducted to undertake significantly larger cohort studies in order to achieve adequate power. Despite our small sample size, our study shows intriguing associations, most likely due to the enrichment of very severe SCZ patients within our study cohort. To the best of our knowledge, our study is the first of its kind to explore the prevalence of known, clinically well-established CNV syndromes within a SCZ patient population. This is highly suggestive that a replication of this study in a much larger sample size with additional indepth phenotyping could show greater enrichment of syndromic CNVs, and further elucidate associations between CNVs and SCZ.

The overall prevalence of syndromic CNVs in SCZ is unknown. In this study, the prevalence of known syndromic CNVs in our SCZ cohort (4.02%) is significantly enriched compared to a reported 0.71% (56/7877) prevalence of such CNVs in an unselected population (P<0.001, 95% C.I. 1.69 – 5.93)(Mefford, 2016). Marshall et al. (2017) identified eight significant genetic loci for SCZ, four of which (1p21.1, 2p16.3, 16p11.2, and 22q11.2) are associated with specific CNV syndromes. We identified three of the same syndromic CNV loci (1p21.1, 2p16.3, and 16p11.2) in our cohort but, interestingly, did not identify any patients with the classic DGS deletion, the most well-known CNV associated with SCZ (Jonas et al., 2014).

In our study, there was a significant enrichment of neurodevelopmental phenotypes within the group of SCZ patients found to carry known syndromic CNVs, an unsurprising result, given the majority of known syndromic CNVs are highly associated and most frequently present with developmental delay (DD) and/or ID first (Coe et al., 2014; Mefford et al., 2012). Interestingly, Kirov and colleagues (2014) found that CNVs associated with SCZ are more likely to result in earlier-onset phenotypes, such as DD, ID and ASD. We also found that patients with greater than two types of hallucinations – thus a more severe SCZ phenotype – were five times more likely to have a known CNV syndrome.

In the group of patients with candidate brain-related CNVs, there was an increased prevalence of TD; patients with TD were 3.6 times more likely to harbour a candidate brain CNV. TD is a side effect of antipsychotics characterized by involuntary movements primarily affecting the orofacial regions (Zai et al., 2018). To date, genetic polymorphisms within dopamine and serotonin receptor, monoamine transporter, and drug metabolizing liver enzyme genes have been associated with TD (Zai et al., 2018). Several studies exploring TD in association with *NRXN1, COMT* and *HLA* resulted in no significant findings (Lanning et al., 2017; Lv et al., 2016). The only TD-associated gene that was also found to overlap one of the CNVs in our cohort was *GPHN*, which encodes an organization protein involved in GABA receptor signaling (Inada et al., 2008); however, the patient with this CNV (33) was not reported to have TD. Although there is little evidence seen for the association of TD with the genes within the candidate brain CNVs identified, tissue specificity analysis of the “candidate brain CNV” gene set reveals the greatest expression in the cerebellum (Supplementary Figure S3), a region of the brain that regulates and coordinates motor movements. Interestingly, one study (Arai et al., 1987) on post-mortem brains of TD patients found a significant inflation of neurons in the cerebellar dentate nucleus.

To date, there is no published data directly exploring the differences in phenotypic presentations between SCZ patients with known syndromic CNVs and those without. Forsingdal et al. (2018) evaluated SCZ-related phenotypes in mouse models harboring SCZ-associated CNVs, and reported that 1q21 deletion mice show altered dopaminergic transmission and response to psychostimulants, both of which are postulated to worsen positive SCZ symptoms (e.g. hallucinations). Microdeletion 22q11.2 mice were also found to show heightened response to NMDA antagonists, hinting at molecular disturbances relevant to positive symptoms in SCZ (Forsingdal et al., 2018). These and our study results highlight the potential impact of CNVs on modulating symptom presentation and severity in SCZ. Further studies are warranted to determine the exact biological mechanisms underlying these effects.

Detailed phenotypic review revealed supportive evidence for the syndromic CNV diagnoses buried in the clinical notes that were focused on psychiatric presentation in several patients (Table 1, bold-bordered cytoband boxes and corresponding bolded phenotypes). This suggests that a broader clinical perspective, including more attention to extra-psychiatric findings, could have increased the likelihood of earlier, accurate diagnosis.

The most striking example was the discovery of microdeletion 17p12, affecting the *PMP22* gene, in two study patients. Deletion of *PMP22* is known to result in hereditary neuropathy with liability to pressure palsies (HNPP; OMIM 162500)(van Paassen et al., 2014). Interestingly, one study (Dracheva et al., 2006) found *PMP22* mRNA levels to be differentially expressed in SCZ. Notably, patient 3 and her father had a documented history of peripheral neuropathies, strongly suggesting that HNPP could have been clinically diagnosed if extra-psychiatric findings had been scrutinized.

Other examples include three patients with microdeletions at chromosome 16p11.2, which is known to be associated with highly penetrant obesity (Perrone et al., 2010; Walters et al., 2010), and two male patients with X chromosome duplications disrupting *CNKSR2*, a gene encoding a synaptic protein in which mutations have been reported in neurodevelopmental disorders and epilepsy (OMIM 301008)(Aypar et al., 2015; Houge et al., 2011; Vaags et al., 2014).

Additional patients were diagnosed with syndromic CNVs that should be easily distinguishable based on clinical findings, but medical records indicate that systemic features were either not ascertained or only limited information was recorded (Table 1, dotted-bordered cytoband boxes).

Patient 10 was found to have a microdeletion at chromosome 5q35.3 encompassing the *NSD1* gene, which is consistent with a diagnosis of Sotos syndrome (OMIM 117550)(Kurotaki et al., 2002). The vast majority of patients with Sotos syndrome (90%) present with the classic triad phenotype of tall stature in childhood with macrocephaly, distinct facial features and ID or learning disability, suggesting that our study patient could likely have been recognized to have Sotos syndrome based on clinical findings (Tatton-Brown et al., 2005; Tatton-Brown and Rahman, 2007).

Patient 13 was found to carry a microdeletion of chromosome 16p13.3 encompassing the *TSC2* and *PKD1* genes, which is a known contiguous gene deletion syndrome comprising phenotypic findings of autosomal dominant tuberous sclerosis (TSC; OMIM 613254) and polycystic kidney disease (PKD; OMIM 173900)(Brook-Carter et al., 1994; Martignoni et al., 2002). Over 90% of TSC patients experience neuropsychiatric and neurodevelopmental disorders in their lifetimes (De Vries et al., 2015). Importantly, targeted treatment of tumours associated with TSC is available, and screening for renal cysts and function in PKD can guide management,(Krueger and Northrup, 2013) highlighting the importance of identifying a genetic diagnosis in psychiatric patients.

Many patients in our cohort were identified to have CNVs containing genes that are known or strong candidates to be associated with SCZ (Table 1, bolded genes).

Four patients were found to have CNVs in 16p13.11. The microdeletion 16p13.11 syndrome has been associated with DD/ID and congenital abnormalities, while duplications in this region have been strongly associated with SCZ (Hannes et al., 2009; Ingason et al., 2011; Ramalingam et al., 2011). Two patients have deletions encompassing only the *ABCC6* gene. While it is currently unknown if deletion of *ABCC6* alone is sufficient to manifest the full microdeletion syndrome, it is of interest that patient 5 has ID/DD, comorbid depression, treatment resistance, and abnormal brain imaging findings. A recent study reported that heterozygous *ABCC6* variants are a risk factor for ischemic stroke and ABCC6 has previously been shown to be enriched in brain microvessel endothelial cells at the blood-brain barrier (BBB)(De Vilder et al., 2018; Warren et al., 2009). Emerging evidence supports that disruptions at the BBB and to brain microvasculature contribute to the development of multiple psychiatric disorders, including SCZ (Kealy et al., 2018). The role of the ATP-binding cassette transporter family, to which ABCC6 belongs, in BBB drug efflux may also suggest a role for ABCC6 variants in treatment response in SCZ and other psychiatric disorders.

Although DGS is the most commonly occurring CNV in SCZ, there were no subjects carrying a typical 3 Mb, or nested 1.5 or 2 Mb DGS deletion in our cohort. Rather, five patients were identified to carry atypical nested deletions at chromosome 22q11.21 encompassing the catechol-O-methyl transferase (*COMT*) gene, two of whom (1 and 2) also carried concomitant known pathogenic CNVs. Due to its role in the metabolism of dopamine, *COMT* is a candidate gene for the development of SCZ (Bassett et al., 2007; Gothelf et al., 2014; Shifman et al., 2002). Multiple studies (Bassett et al., 2007; Gothelf et al., 2005; Gothelf et al., 2014; Murphy et al., 1999) have shown conflicting results of whether a less functional *COMT* allele increases the risk of psychosis. The enrichment of this atypical deletion within our cohort may provide additional support for the importance of *COMT* in the development of SCZ.

Perhaps not surprisingly, most of the SCZ-related genes seen within the CNVs identified have also been associated with other neuropsychiatric and neurodevelopmental conditions, contributing further evidence to support the neurodevelopmental continuum model (Davis et al., 2016; Owen and O’Donovan, 2017). For example, the *RMB8A* gene contained within the 1q21.1 microdeletion in patient 15 has been associated with ID and anxiety behaviours (Alachkar et al., 2013; Gamba et al., 2016; Zou et al., 2015), while deletions of *NRXN1*, seen in two patients, are strongly associated with neuropsychiatric and neurodevelopmental disorders, including SCZ and ASD (Kirov et al., 2009; Marshall et al., 2017; Rujescu et al., 2008). Deletion of *LMAN2L*, a gene associated with increased risk for SCZ, bipolar disorder and ID (Lim et al., 2014; Rafiullah et al., 2016), was found in three patients. One of the most frequent deletions found was at chromosome 5p12, involving the *HCN1* gene, which encodes a voltage-gated potassium channel, mutations in which cause early infantile epileptic encephalopathy (Nolan et al., 2004; Nolan et al., 2003; Santoro et al., 2010). It is postulated that disrupted synaptic transmission seen in HCN1-deficient humans and mice could be playing a role, not just in epilepsy, but also in memory formation and learning (Nava et al., 2014; Nolan et al., 2004; Santoro et al., 2010). Indeed, individuals with *HCN1* mutations have been reported with ID and autistic features (Nava et al., 2014). Emerging evidence supporting a fluid spectrum and common underlying genetic mechanisms for many neurodevelopmental, neurological and psychiatric disorders prompts the need for further study of genes currently primarily associated with only one of these realms (Anttila et al., 2018).

Across all CNVs identified in our study cohort, gene pathway enrichment analysis identified enrichment of immune system pathways, particularly the adaptive immune system and cytokine signaling pathways. Furthermore, patients with CNVs containing immune system genes were found to have higher total number of CNVs, average number of brain-related CNVs and proportion of brain CNVs to total CNV number. These results suggest that brain- and immune-related CNVs in SCZ patients are correlated with one another and travel together genetically. The enrichment of immune pathways supports decades of research showing a strong relationship between immune genes and environmental factors in the development of SCZ (Cattane et al., 2018).

The immune hypothesis of SCZ was borne by an epidemiological study in Finland where Mednick et al. (1988) found that offspring of pregnant mothers during the 1957 influenza epidemic had a higher chance of developing SCZ. To date, this hypothesis has been supported by numerous genetic studies, including the finding of the Major Histocompatibility (MHC) region being the strongest signal in genome-wide association studies of SCZ (Schizophrenia Working Group of the Psychiatric Genomics Consortium, 2014; Shi et al., 2009; Stefansson et al., 2009). Our study identified one patient (32) with a CNV in the MHC region, overlapping genes of interest *HLA-B* and *MICB*. Polymorphisms in the *MICB* gene have been associated with grey matter volume and working memory, while *HLA-B* has been implicated in the life cycles of pathogens in SCZ patients (Carter, 2009; Shirts et al., 2007).

Taken together, our results support the hypothesis that syndromic CNVs are enriched in SCZ patient populations, and provide further evidence that CNVs in general play a significant role in SCZ risk and, potentially, in modulating SCZ sub-phenotypes. Based on our regression analysis, we propose a CNV-SCZ model (Figure 2) in which CNVs could be effecting the clinical presentations of SCZ patients. While neurodevelopmental presentations are most often the earliest clinical symptoms seen in individuals with known syndromic CNVs, the presence of both syndromic and immune-related CNVs could potentially increase the risk for development of SCZ through external, environmental interactions. Furthermore, the combination of syndromic and immune-related CNVs could potentiate exacerbated symptomatology in patients diagnosed with SCZ. Based on our proposed model, we recommend that SCZ patients with a phenotypic profile consisting of neurodevelopmental presentations alone or along with greater than two types of hallucinations should undergo CNV analysis to identify potential underlying syndromic CNVs.

**Figure 2.**
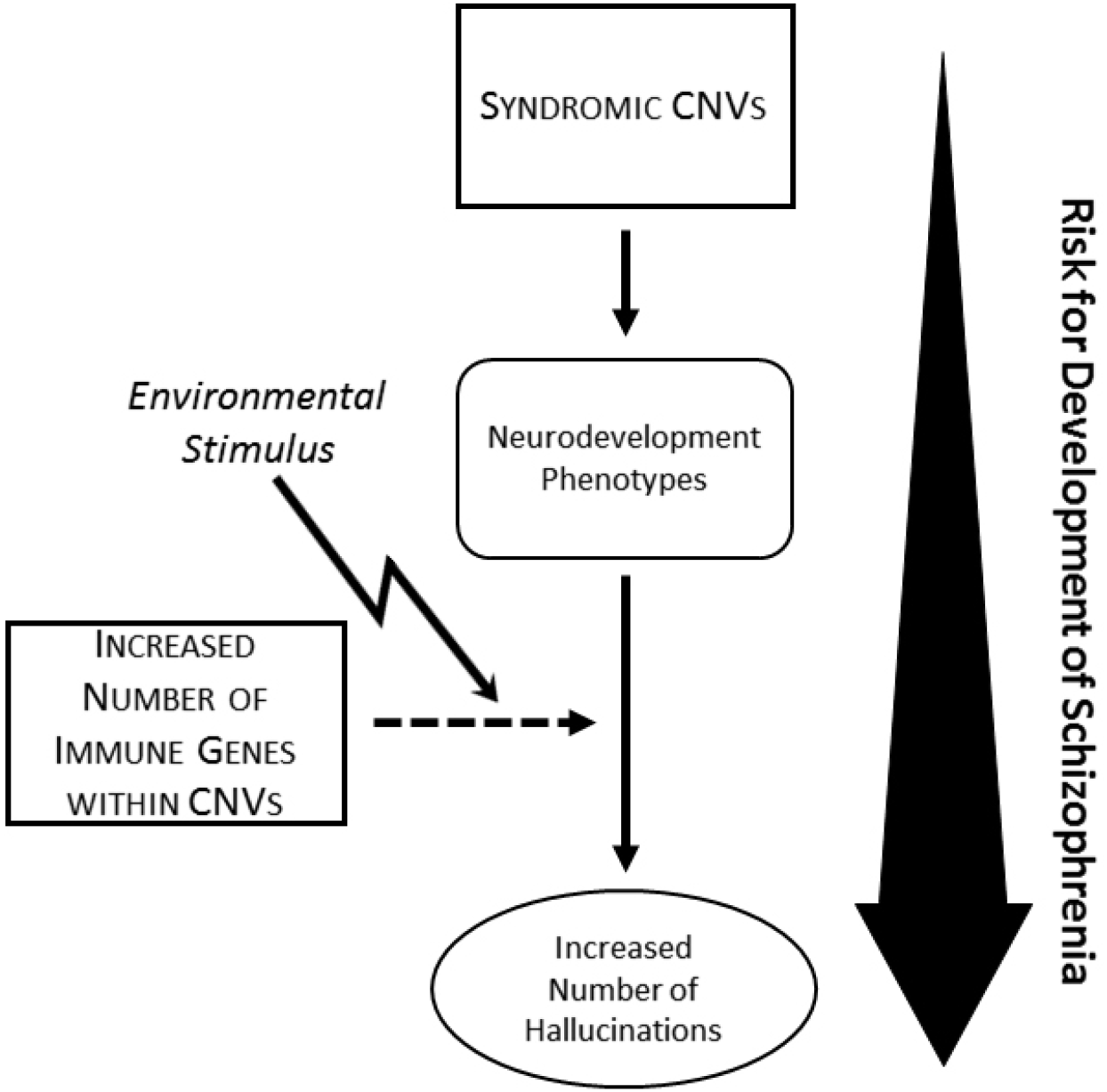
Proposed model of CNV syndromes in SCZ. Syndromic CNVs lead to neurodevelopmental phenotypes first. Those with greater numbers of immune-related CNVs are at increased risk of developing SCZ through interaction with external, environmental stimulus and increased severity of psychotic symptoms.

