## Supplementary Materials for "Copy number variant syndromes are frequent in schizophrenia: progressing towards a CNV-schizophrenia model"

**Supllementary Tables**

**Table 1.** Demographic characteristics of the study sample (n=348)*

| **Characteristic** | **Average Age (S.D)** | **Male (%) : Female (%)** | **Ethnicity (%)** | | | | |
| --- | --- | --- | --- | --- | --- | --- | --- |
|  |  |  | ***Caucasian*** | ***African*** | ***East Asian*** | ***South Asian*** | ***Other*** |
|  | 49.8 (11.8) | 240 (69) : 108 (31) | 267 (77) | 38 (11) | 26 (8) | 5 (1) | 12 (3) |

**This sample set has been quality controlled for relatedness amongst patients and any patients with >0.25 relatedness were removed prior to analyses.* S.d., standard deviation.

**Table S2.** Gene enrichment pathway analyses results from GO and Reactome.

| **Enriched Pathways** | **P_FDR_** |
| --- | --- |
| Alpha-defensins (R-HSA-1462054) | 3.99E-02 |
| Phase II conjugation (R-HSA-156580) | 4.10E-02 |
| Immunoregulatory interactions between a Lymphoid and a non-Lymphoid cell (R-HSA-198933) | 2.80E-02 |
| Immune System (R-HSA-168256) | 4.43E-02 |

| **ID**  **Table S3.** Patient CNV and phenotype data. | **Cytoband** | **Genomic Coordinates (GRCh37/hg19)** | **Genes** |
| --- | --- | --- | --- |
| 15 | **Del 1q21.1** | chr1:144723763-146297807 | HFE2A*, RBM8A,* ***PEX11B****,* NBPF9, SEC22L, PIAS3, POLR3C, CD160, PDZK1 |
| 1 | **Del 1q21.1-q21.2** | chr1:146780414-147526040 | *GJA5,* ***GJA8****,* BCL9, GPR89B |
|  | **Del 16p11.2** | chr16:29474810-30099408 | KIF22, PRRT2*,* MAZ |
|  | Del 22q11.21 | chr22:19925029-20175355 | ***COMT****, TANGO2,* ARVCF, MIR185, DGCR8, RANBP1, ZDHHC8 |
| 21 | **Del 2p16.3** | chr2:50631006-50870834 | ***NRXN1*** |
| 22 | **Del 2p16.3** | chr2:50267675-50479696 | ***NRXN1*** |
|  | Del 2p16.3 | chr2:49167207-49394694 | *FSHR* |
| 10 | **Del 5q35.3** | chr5:176662203-176879182 | ***NSD1****,* SLC34A1, F12 |
|  | Del 19p13.3 | chr19:938571-1231659 | STK11*,* ABCA7 |
| 11 | **Del 15q11.1-q12** | chr15:20224763-26500067 | *NIPA1,* MKRN3*,* MAGEL2*, NDN, SNRPN, UBE3A,* TUBGCP5, CYFIP1, NIPA2 |
| 12 | **Del 16p11.2** | chr16:29648146-31122248 | KIF22, PRRT2*,* ***ALDOA****,* **TBX6**, CORO1A, SRCAP, PHKG2, HSD3B7, STX1B, VKORC1, BCKDK*,* MAZ, **SEZ6L2**, **HIRIP3**, **DOC2A**, MAPK3 |
|  | Del 5q23.3-q31.1 | chr5:130533005-130767904 | LYRM7*,* RAPGEF6 |
|  | Del 11q13.1 | chr11:65358334-65624227 | RNASEH2C*,* MAP3K11, NFKB3, KAT5, CFL1 |
|  | Del 19p13.2-p13.12 | chr19:13721364-14102566 | *CC2D1A,* NOS3, MIR181C |
| 13 | **Del 16p11.2** | chr16:29626237-31184275 | KIF22, PRRT2*,* ***ALDOA****,* **TBX6**, CORO1A, SRCAP, *PHKG2,* HSD3B7, STX1B, VKORC1, BCKDK, MAZ, **SEZ6L2**, **HIRIP3**, **DOC2A**, MAPK3 |
|  | **Del 16p13.3** | chr16:1898885-2196820 | *GFER,* NTHL1, ***TSC2****,* PKD1*, SYNGR3* |
|  | Del 11q13.1 | chr11:64230769-65193674 | SLC22A12, RASGRP2, PYGM, MEN1, CAPN1, NRXN2 |
|  | Del 19p13.3 | chr19:388821-2279433 | BSG*,* ELANE, CFD, KISS1R, ABCA7, GPX4, STK11*, NDUFS7, GAMT,* APC2, REEP6, TCF3, *ADAT3,* AP3D1, AMH, MADCAM1, CDC34, GZMM, POLRMT, FGF22, FSTL3, PALM, PRTN3 |
|  | Del 19p13.2-p13.12 | chr19:13703822-14596500 | *CC2D1A,* PRKACA*,* NOS3, MIR181C |
|  | Del 19q13.33 | chr19:50617963-51018893 | MYH14, *KCNC3,* POLD1, NR1H2 |
|  | Del 19q13.42 | chr19:54648737-55238193 | *MBOAT7, TSEN34,* CNOT3, RPS9, LILRB3, LILRA3, CDC42EP5, LILRA1 |
| 4 | **Del 16p13.11-p12.3** | chr16:15389435-18089485 | *NDE1, MYH11,* ***ABCC6****, XYLT1,* FOPNL, ABCC1 |
| 14 | **Dup 16p13.11-p12.3** | chr16:15389435-18154791 | *NDE1, MYH11,* ***ABCC6****, XYLT1,* ABCC1 |
| 2 | **Del 17p12** | chr17:14023683-15421887 | *COX10,* ***PMP22****,* TEKT3 |
|  | Del 22q11.21 | chr22:19795026-20038108 | ***COMT****, TANGO2,* GNB1L, ARVCF, MIR185 |
| 3 | **Del 17p12** | chr17:14034987-15425596 | *COX10,* ***PMP22*** |
| 37 | **Dup Xp22.12** | chrX:20923213-21580432 | ***CNKSR2*** |
|  | Dup 19q13.2 | chr19:41705425-42677868 | *TGFB1,* B9D2, BCKDHA, RPS19, CD79A, ATP1A3 |
|  | Dup Xp22.11 | chrX:23583405-24370505 | *KLHL15,* EIF2S3 |
|  | Dup Xp11.22 | chrX:54097999-54632588 | TSR2, FGD1, FAM120C |
| 44^†^ | **Dup Xp22.12** | chrX:20923213-21580432 | ***CNKSR2*** |
|  | Dup 2q33.1-q33.1 | chr2:202770323-204243381 | SUMO1, BMPR2, ABI2 |
|  | Dup 4p14 | chr4:39225506-39768600 | WDR19, LIAS, UBE2K |
|  | Dup 10q23.32-q23.33 | chr10:69238823-70255706 | *CTNNA3*, DNAJC12, MYPN, ATOH7, DNA2 |
| **ID** | **Cytoband** | **Genomic Coordinates (GRCh37/hg19)** | **Genes** |
| 24 | Del 1q43 | chr1:237416669-237631305 | *RYR2* |
| 31 | Del 2p16.3 | chr2:49121934-49354502 | *FSHR* |
|  | Del 19p12-q13.11 | chr19:24194082-32779217 | *C19orf12,* UQCRFS1, TSHZ3 |
| 35 | Del 2p22.3 | chr2:32481501-33185282 | *NLRC4* |
| 23 | Del 2q11.2 | chr2:97108075-97429511 | ***LMAN2L****,* CNNM4 |
| 40 | Del 2q11.2 | chr2:97108075-97429511 | ***LMAN2L****,* CNNM4 |
| 42 | Del 2q11.2 | chr2:97108075-97429511 | ***LMAN2L****,* CNNM4 |
| 36 | Del 2q21.2 | chr2:131194418-131701010 | CFC1, ARHGEF4 |
| 19 | Del 3p26.3-p26.2 | chr3:2757210-3182640 | TRNT1, **CNTN4** |
| 32 | Del 3p26.2 | chr3:3161894-3451929 | TRNT1, *CRBN* |
|  | Del 6p21.33 | chr6:30800518-31473848 | VARS2, CDSN, *HLA-C,* ***HLA-B,* MICB** |
|  | Del 14q32.33 | chr14:105737185-106031276 | *BRF1,* MTA1, CRIP2 |
| 39 | Del 5p12 | chr5:45209896-45715271 | ***HCN1*** |
| 33 | Del 5p12 | chr5:45284212-45752474 | ***HCN1*** |
|  | Del 14q23.3 | chr14:67477553-67702607 | *GPHN* |
| 41 | Del 5p12 | chr5:45372258-45715271 | ***HCN1*** |
|  | Del 19p12-q13.11 | chr19:24164923-32512819 | *C19orf12,* UQCRFS1, TSHZ3 |
| 30 | Del 5p12 | chr5:45491554-45715271 | ***HCN1*** |
|  | Del 16p13.3 | chr16:1408394-1937615 | *GNPTG,* CLCN7, *TELO2,* IFT140, *MRPS34,* IGFALS, HAGH, FAHD1 |
| 29 | Del 5p12 | chr5:45510921-45914084 | ***HCN1*** |
|  | Del 15q15.3-q21.1 | chr15:44795340-45093008 | *SPG11,* B2M |
| 43 | Del 5q14.2 | chr5:82432502-82645969 | *XRCC4* |
| 26 | Del 6q26 | chr6:162550689-162988063 | *PRKN* |
| 20 | Del 9p24.3 | chr9:372245-723212 | DOCK8, KANK1 |
| 25 | Del 10q11.21-q11.22 | chr10:45905767-47241036 | *ALOX5,* SYT15, GPRIN2, NPY4R |
| 27 | Del 10q11.22 | chr10:48350704-48882834 | RBP3, GDF2, GDF10 |
| 34 | Del 10q21.3 | chr10:67908524-68112828 | *CTNNA3* |
| 17 | Del 11q13.2 | chr11:67239223-67505898 | AIP, CABP2, NDUFV1, PITPNM1 |
| 38 | Del 15q14 | chr15:34585121-34954567 | SLC12A6, NOP10, GOLGA8A, GOLGA8B |
| 5 | Del 16p13.11 | chr16:16237991-16588399 | ***ABCC6*** |
| 6 | Del 16p13.11 | chr16:16237991-16706164 | ***ABCC6*** |
| 18 | Del 17p13.1 | chr17:8699348-9839794 | *PIK3R5,* PIK3R6, DHRS7C, GAS7 |
| 28 | Del 19p12-q13.11 | chr19:24172349-32657355 | *C19orf12,* UQCRFS1, TSHZ3 |
| 7 | Del 22q11.21 | chr22:19872615-20175355 | ***COMT****, TANGO2,* ARVCF, MIR185, DGCR8, RANBP1, ZDHHC8 |
| 8 | Del 22q11.21 | chr22:19872615-20175355 | ***COMT****, TANGO2,* ARVCF, MIR185, DGCR8, RANBP1, ZDHHC8 |
| 9 | Del 22q11.21 | chr22:19925029-20175355 | ***COMT****, TANGO2,* ARVCF, MIR185, DGCR8, RANBP1, ZDHHC8 |
| 16 | Dup Xq26 | chrX:154235666-154887040 | F8, *RAB39B, CLIC2,* TMLHE, SPRY3 |

Bolded cytobands are those of known syndromic CNVs. All regular font genes listed are OMIM genes associated with a particular disease phenotype, italicized genes are OMIM genes associated with neuronal/psychiatric phenotypes, underlined genes are non-OMIM genes associated with neuronal/psychiatric phenotypes, and bolded genes are specifically associated with schizophrenia. All phenotype presentations characteristic of their respective CNV syndromes are bolded. All listed genes within duplications are disrupted by the duplication. †Subject 44 did not have any phenotype information available. DSM-V, Diagnostic and Statistical Manual of Mental Disorders-5; DD, developmental delay; ID, intellectual disability; Hall, hallucination; TRx, treatment resistance; Neuro, neurological phenotypes; Hx, history; psych, psychiatric; FHx, family history; DEP, depressive disorders; PD, personality disorders; SWD, sleep-wake disorders; ADHD, attention deficit hyperactive disorder; ED, elimination disorders; ANX, anxiety disorders; SS, schizophrenia spectrum and other psychotic disorders; BPD, bipolar and related disorders; CD, disruptive, impulse-control, and conduct disorders; SD, somatic symptoms and related disorders; TD, trauma – and stressor – related disorders; MOT, motor disorders; OCD, obsessive-compulsive and related disorders; A, auditory hallucination; V, visual hallucination; T, tactile hallucination; G/O, gustatory and/or olfactory hallucination; Ep, epilepsy; Ab, abnormal; MS, multiple sclerosis; T2DM, type 2 diabetes mellitus; DM, diabetes mellitus; O, obesity; HN, hernia (abdnomina, inguinal); SIDS, sudden infant death syndrome; LD, learning disability; T21, trisomy 21. Obesity was defined as a BMI≥30 or patient medical chart notes that the subject was obese.

| **Gene**  **Table S4.** Complete list of “candidate brain” genes and relevant literature. | **Gene cards ID** | **Associated phenotype** | **Relevant Literature** |
| --- | --- | --- | --- |
| *RYR2* | GC01P237042 | SCZ, ID | Basset *et al.* (2017)^1^; Hamdan *et al.* (2014)^2^; Ambalavalan *et al.* (2016)^3^ |
| *FSHR* | GC02M048866 | SCZ, BPD, MDD, AD | Chen *et al.* (2015)^10^; Corbo *et al.* (2011)^11^; Sun *et al.* (2014)^12^ |
| *C19orf12* | GC19M029699 | neuropsychiatric, neurodegeneration | Klysz *et al.* (2014)^13^; Heidari *et al.* (2016)^14^; Paudel *et al.* (2015)^15^; Aoun and Tiranti (2015)^16^ |
| *UQCRFS1* | GC19M029205 | SCZ, SAF | Takao *et al.* (2013)^17^; Arion *et al.* (2015)^18^ |
| *TSHZ3* | GC19M031274 | ASD | Caubit *et al.* (2016)^19^ |
| *NLRC4* | GC02M032224 | neurodegeneration | Freeman and Ting (2016)^20^ |
| *LMAN2L* | GC02M096793 | SCZ, BPD, ID | Lim *et al.* (2014)^21^; Khan *et al.* (2016)^22^; Rafiullah *et al.* (2016)^23^ |
| *ARHGEF4* | GC02P130836 | ADHD, DD, ID, Ep, neurobehavioral, ASD | Dharmadhikari *et al.* (2012)^24^; Eriksson *et al.* (2015)^25^ |
| *ABI2* | GC02P203327 | ASD, ID | Grove *et al.* (2004)^26^; Lee *et al.* (2015)^27^; Hlushchenko *et al.* (2016)^28^; Guo *et al.* (2017)^29^; Bardoni and Abekhoukh (2014)^30^; Harripaul *et al.* (2018)^31^; Durand *et al.* (2012)^32^ |
| *UBE2K* | GC04P039700 | PD, neurological | Ryan *et al.* (2006)^33^; Molochnikov *et al.* (2012)^34^; Kaytor and Warren (1999)^35^; Anuppalle *et al.* (2013)^36^ |
| *CTNNA3* | GC10M065912 | Ep, ASD, AD, DD, TS | Lesca *et al.* (2012)^37^; Lintas *et al.* (2017)^38^; Miyashita *et al.* (2007)^39^; Allen *et al.* (2015)^40^; Armour *et al.* (2016)^41^; Shimojima *et al.* (2018) ^42^; Fernandez (2016)^43^; Fang *et al.* (2017)^44^; Chen *et al.* (2013)^45^; Bachelli *et al.* (2014)^46^ |
| *CNTN4* | GC03P002117 | SCZ, ASD, MR | Burbach and va der Zwaag (2009)^52^; Molenhuis *et al.* (2016)^53^; Zhao *et al.* (2013)^54^; Dijkhuizen *et al.* (2006)^55^; Cottrell *et al.* (2011)^56^; Boraska *et al.* (2014)^57^; Goes *et al.* (2015)^58^ |
| *CRBN* | GC03M003166 | MR | Dijkhuizen *et al.* (2006)^55^; Xin *et al.* (2008)^59^ |
| *HLA-C* | GC06M031272 | SZ, MS | ISGC (2012)^60^; Andreassen *et al.* (2015)^61^ |
| *HLA-B* | GC06M031277 | SZ, MS | ISGC (2012)^60^; Andreassen *et al.* (2015)^61^; Palmer *et al.* (2006)^62^ |
| *BRF1* | GC14M105212 | ID, neurodevelopmental | Nevado *et al.* (2014)^63^; Brock *et al.* (2015)^64^ |
| *MTA1* | GC14P105419 | ID, neurodevelopmental | Nevado *et al.* (2014)^63^ |
| *CRIP2* | GC14P105472 | ID, neurodevelopmental | Nevado *et al.* (2014)^63^ |
| *HCN1* | GC05M045260 | SCZ, AD, BPD, SZ | Neymotin *et al.* (2016)^65^; Berridge (2013)^66^; Nolan *et al.* (2004)^67^; Nolan *et al.* (2003)^68^; Santoro *et al.* (2010)^69^ |
| *GPHN* | GC14P066507 | SCZ, ASD, SZ, Ep, neurodevelopmental | Lionel *et al.* (2013)^70^; Hu *et al.* (2015)^71^; Dejanovic *et al.* (2014)^72^ |
| *GNPTG* | GC16P001351 | SCZ, Dys, ID, ASD, Ep, BPD, neurodevelopmental | Zhao *et al.* (2015)^73^; Chen *et al.* (2015)^74^; Kang *et al.* (2015)^75^; Deriziotis and Fisher (2017)^76^; Kazemi *et al.* (2017)^77^; Ewald *et al.* (2002)^78^; Lourov *et al.* (2012)^79^ |
| *TELO2* | GC16P001493 | ID, ASD, Ep, BPD | Ewald *et al.* (2002)^78^; Lourov *et al.* (2012)^79^ |
| *MRPS34* | GC16M001771 | ID, ASD, Ep, BPD | Ewald *et al.* (2002)^78^; Lourov *et al.* (2012)^79^ |
| *FAHD1* | GC16P001826 | ID, ASD, Ep, BPD | Ewald *et al.* (2002)^78^; Lourov *et al.* (2012)^79^; Hashimoto *et al.* (2016)^80^ |
| *SPG11* | GC15M044562 | neurodevelopmental | Mishra *et al.* (2016)^81^; Mishra (2014)^82^ |
| *XRCC4* | GC05P083077 | SCZ | Wang *et al.* (2010)^83^; Pehlivan *et al.* (2017)^84^; Mazaheri and Saadat (2015)^85^ |
| *PRKN* | GC06M161348 | PD | Hedrich *et al.* (2006)^86^; Pramstaller *et al.* (2005)^87^; Mellick *et al.* (2009)^88^ |
| *DOCK8* | GC09P000214 | MR, DD, ID, ASD | Tassano *et al.* (2016)^89^; Wang *et al.* (2016)^90^ |
| *KANK1* | GC09P000474 | MR, DD, ID, ASD | Tassano *et al.* (2016)^89^; Wang *et al.* (2016)^90^ |
| *ALOX5* | GC10P045338 | AD, SCZ | Šerý *et al.* (2016)^91^; Tang *et al.* (2012)^92^; Grayson *et al.* (2013)^93^ |
| *SYT15* | GC10P046578 | PD | La Cognata *et at.* (2017)^94^ |
| *GPRIN2* | GC10M046543 | SCZ, BPD | Nuttle (2016)^95^; Ghai *et al.* (2011)^96^; Chen *et al.* (2016)^97^ |
| *NPY4R* | GC10M046461 | SCZ, ID | Rodriguez-Santiago *et al.* (2010)^98^; Qiao *et al.* (2010)^99^ |
| *GDF10* | GC10P047300 | ASD, neurological | Jie (2004)^100^; Li *et al.* (2015)^101^; Caubit *et al.* (2016)^19^; Carmicheal (2016)^102^; Carmicheal *et al.* (2016)^103^; Li *et al.* (2010)^104^; Kashima and Kata (2017)^105^ |
| *PITPNM1* | GC11M067492 | SCZ, ASD, ID | McCarthy *et al.* (2014)^106^; Alhuzimi *et al.* (2018)^107^ |
| *GOLGA8A* | GC15M034380 | SCZ, ID | Nevado *et al.* (2014)^63^; Freedman and Leonard (2001)^108^; Antonacci *et al.* (2014)^109^ |
| *GOLGA8B* | GC15M034525 | SCZ, ID | Nevado *et al.* (2014)^63^; Freedman and Leonard (2001)^108^; Antonacci *et al.* (2014)^109^ |
| *PIK3R5* | GC17M008878 | ID, SCZ, ADHD | Nevado *et al.* (2014)^63^; Gross and Bassell (2014)^110^; Lesch *et al.* (2008)^111^ |
| *PIK3R6* | GC17M008802 | ID, SCZ, ADHD | Nevado *et al.* (2014)^63^; Gross and Bassell (2014)^110^; Lesch *et al.* (2008)^111^ |
| *DHRS7C* | GC17M009771 | ID, SCZ, ADHD | Nevado *et al.* (2014)^63^; Gross and Bassell (2014)^110^; Lesch *et al.* (2008)^111^ |
| *GAS7* | GC17M009910 | ID, SCZ, ADHD | Nevado *et al.* (2014)^63^; Gross and Bassell (2014)^110^; Lesch *et al.* (2008)^111^; Zhang *et al.* (2016)^112^ |
| *TGFB1* | GC19M041301 | ID, SZ, SCZ, DD, Ob | Nevado *et al.* (2014)^63^; Hall *et al.* (2010)^113^; Tayeh *et al.* (2015)^114^; Rim *et al.* (2017)^115^; Nacinovich *et al.* (2017)^116^ |
| *KLHL15* | GC0XM023911 | MR | Mignon-Ravix *et al.* (2014)^117^; AlSagob *et al*. (2015)^118^ |
| *FAM120C* | GC0XM054097 | ASD, ID | De Wolf *et al*. (2014)^119^; Santos- Rebouças *et al*. (2015)^120^, Qiao *et al*. (2008)^121^; Edens *et al*. (2011)^122^ |
| *RAB39B* | GC0XM155259 | ID, DD | Vanmarsenille *et al*. (2014)^123^; Anderson *et al*. (2014)^124^; El-Hattab *et al*. (2015)^125^; El-Hattab *et al*. (2011)^126^; El-Hattab *et al*. (2016)^127^ |
| *CLIC2* | GC0XM155276 | ID, DD | El-Hattab *et al*. (2016)^127^; Anderson *et al*. (2014)^124^; El-Hattab *et al*. (2015)^125^; El-Hattab *et al*. (2011)^126^ |
| *SPRY3* | GC0XP155612 | ASD | Ning *et al.* (2015)^128^ |

Literature search was performed using PubMed, OMIM, GeneCards, and Google Scholar. SCZ, schizophrenia; ID, intellectual disability; ASD, autism spectrum disorder; BPD, bipolar disorder; MDD, major depressive disorder; AD, Alzheimer’s disorder; SAF, schizoaffective disorder; ADHD, attention deficit hyperactive disorder; DD, developmental delay; Ep, epilepsy; PD, Parkinson’s disease; TS, Tourette’s syndrome; SZ, seizures; MR, mental retardation; MS, multiple sclerosis; Dys, dyslexia; Ob, obesity.

**Table S5.** *Post-hoc* power analysis.

| **Statistical Test** | **Group 1** | **Group 2** | **Achieved Power/MDE** |
| --- | --- | --- | --- |
| Mann-Whitney-U | Known | None | Power: 0.99 |
|  | Known | Candidate | Power: 0.07 |
|  | Candidate | None | Power: 1.00 |
| Chi-Square Test | Phenotypic Comparisons | | MDE: 0.15 |
| Exact Binomial | Sample Known Syndromic CNV Prevalence | Mefford, 2016 Prevalence | Power: 0.987 |
|  | Sample Prevalence | Known Population Prevalence | MDE: 0.02 |

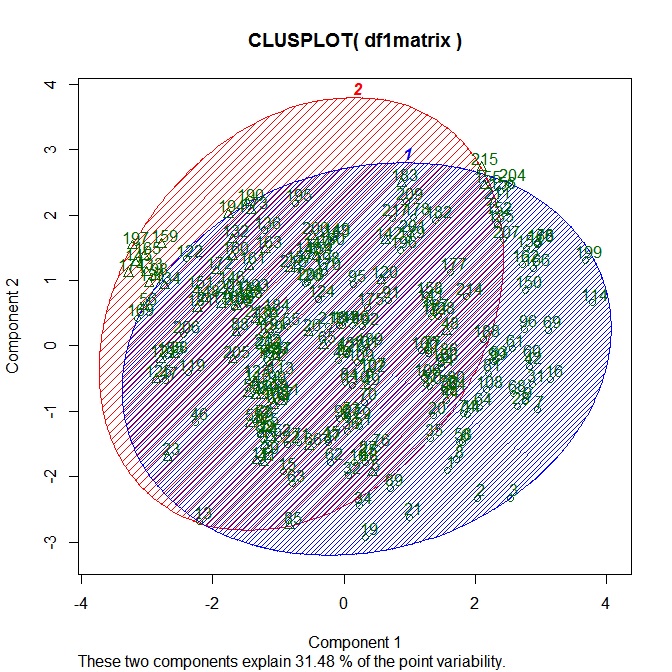
**Supplementary Figures**

**Figure S1.** K-modes clustering graphical output for k=2. Briefly, fviz_nbclust code from R package ‘factoextra’ was used to generate a silhouette plot in order to determine k, the optimal number of clusters. K-modes clustering analysis was performed using kmodes from ‘klaR v0.6-14’ package and clusplot from ‘cluster v2.0.7-1’ was used to visualize the cluster analysis.

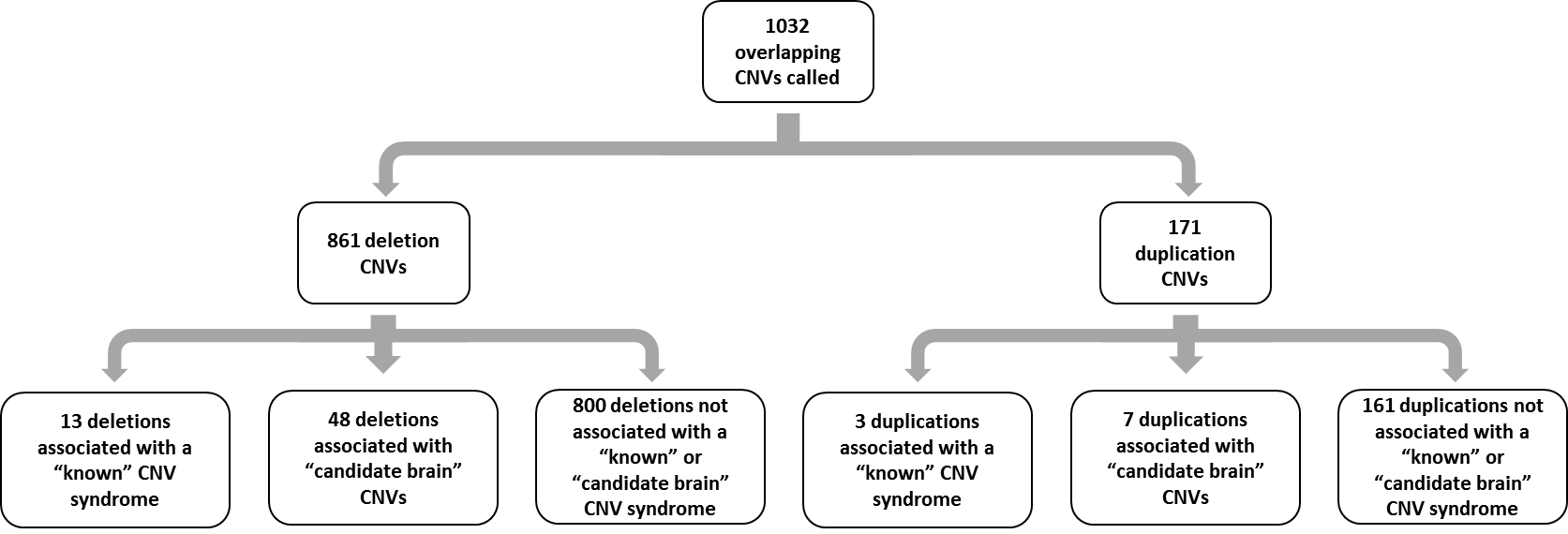

**Figure S2.** Breakdown of all CNVs by group. (a) CNV categorization based on presence (“immune”) or absence (“non-immune”) of immune pathway genes by total number of patients. (b) CNV breakdown based on tissue specificity analyses by copy number and CNV groups ("known", "candidate brain" and "none").

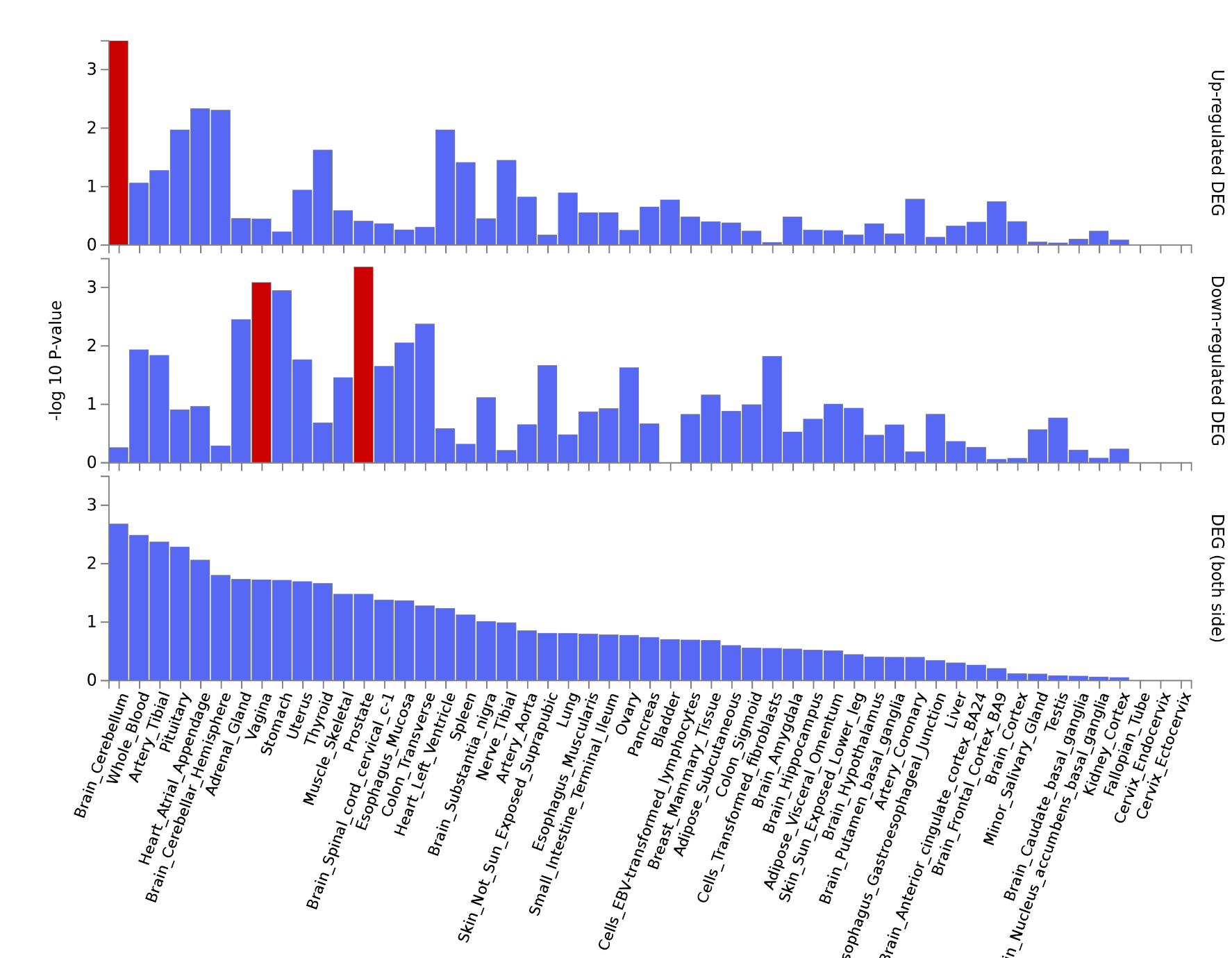

**Figure S3.** Tissue specificity graph of differentially expressed gene (DEG) set only from the “candidate brain CNV” group. Significantly enriched DEG sets (Pbon < 0.05) are depicted by the red bars. Image generated by FUMA GENE2FUNC.
**References**

**1.**	Bassett AS, Lowther C, Merico D, et al. Rare genome-wide copy number variation and expression of schizophrenia in 22q11. 2 deletion syndrome. *American Journal of Psychiatry* 2017;174(11):1054-1063.

**2.**	Hamdan FF, Srour M, Capo-Chichi J-M, et al. De novo mutations in moderate or severe intellectual disability. *PLoS genetics* 2014;10(10):e1004772.

**3.**	Ambalavanan A, Girard SL, Ahn K, et al. De novo variants in sporadic cases of childhood onset schizophrenia. *European Journal of Human Genetics* 2016;24(6):944.

**4.**	Mitchell KJ, Porteous DJ. Rethinking the genetic architecture of schizophrenia. *Psychological medicine* 2011;41(1):19-32.

**5.**	Kasem E, Kurihara T, Tabuchi K. Neurexins and neuropsychiatric disorders. *Neuroscience research* 2017.

**6.**	Wang J, Gong J, Li L, et al. Neurexin gene family variants as risk factors for autism spectrum disorder. *Autism Research* 2018;11(1):37-43.

**7.**	Kirov G, Rujescu D, Ingason A, Collier DA, O'donovan MC, Owen MJ. Neurexin 1 (NRXN1) deletions in schizophrenia. *Schizophrenia bulletin* 2009;35(5):851.

**8.**	Rujescu D, Ingason A, Cichon S, et al. Disruption of the neurexin 1 gene is associated with schizophrenia. *Human molecular genetics* 2008;18(5):988-996.

**9.**	Ching MS, Shen Y, Tan WH, et al. Deletions of NRXN1 (neurexin‐1) predispose to a wide spectrum of developmental disorders. *American Journal of Medical Genetics Part B: Neuropsychiatric Genetics* 2010;153(4):937-947.

**10.**	Chen X, Long F, Cai B, Chen X, Chen G. A novel relationship for schizophrenia, bipolar and major depressive disorder: a hint from chromosome 2 high density association screen. 2015.

**65.**	Neymotin SA, Sherif MA, Jung JQ, Kabariti JJ, Lytton WW. Genome-wide associations of schizophrenia studied with computer simulation. 2016.

**77.**	Kazemi N, Estiar MA, Fazilaty H, Sakhinia E. Variants in GNPTAB, GNPTG and NAGPA genes are associated with stutterers. *Gene* 2017.

**78.**	Ewald H, Flint T, Kruse T, Mors O. A genome-wide scan shows significant linkage between bipolar disorder and chromosome 12q24. 3 and suggestive linkage to chromosomes 1p22–21, 4p16, 6q14–22, 10q26 and 16p13. 3. *Molecular Psychiatry* 2002;7(7):734.

**79.**	Iourov IY, Vorsanova SG, Kurinnaia OS, Zelenova MA, Silvanovich AP, Yurov YB. Molecular karyotyping by array CGH in a Russian cohort of children with intellectual disability, autism, epilepsy and congenital anomalies. *Molecular Cytogenetics* 2012;5(1):46.

**80.**	Hashimoto R, Nakazawa T, Tsurusaki Y, et al. Whole-exome sequencing and neurite outgrowth analysis in autism spectrum disorder. *Journal of human genetics* 2016;61(3):199.

**81.**	Mishra HK, Prots I, Havlicek S, et al. GSK3ß‐dependent dysregulation of neurodevelopment in SPG11‐patient induced pluripotent stem cell model. *Annals of neurology* 2016;79(5):826-840.

**82.**	Mishra HK. Modeling neurodevelopment and cortical dysfunction in SPG11-linked hereditary spastic paraplegia using human induced pluripotent stem cells.

**95.**	Nuttle A. *Human-specific duplicate genes: new frontiers for disease and evolution*; 2016.

**96.**	Ghai SJ, Shago M, Shroff M, Yoon G. Cockayne syndrome caused by paternally inherited 5Mb deletion of 10q11. 2 and a frameshift mutation of ERCC6. *European journal of medical genetics* 2011;54(3):272-276.

**97.**	Chen J, Calhoun V, Perrone-Bizzozero N, Pearlson G, Sui J, Du Y, Liu J. A pilot study on commonality and specificity of copy number variants in schizophrenia and bipolar disorder. *Translational psychiatry* 2016;6(5):e824.

**98.**	Rodriguez-Santiago B, Brunet A, Sobrino B, et al. Association of common copy number variants at the glutathione S-transferase genes and rare novel genomic changes with schizophrenia. *Molecular psychiatry* 2010;15(10):1023.

**99.**	Qiao Y, Harvard C, Tyson C, et al. Outcome of array CGH analysis for 255 subjects with intellectual disability and search for candidate genes using bioinformatics. *Human genetics* 2010;128(2):179-194.

**100.**	Jie L. Factors regulating hippocampal neuronal proliferation, differentiation and survival. 2004.

**127.**	El-Hattab AW, Schaaf CP, Cheung SW. Xq28 duplication syndrome, Int22h1/Int22h2 mediated. 2016.

**128.**	Ning Z, McLellan AS, Ball M, et al. Regulation of SPRY3 by X chromosome and PAR2-linked promoters in an autism susceptibility region. *Human molecular genetics* 2015;24(18):5126-5141.
